# HYD3, a conidial hydrophobin of the fungal entomopathogen *Metarhizium acridum* induces the immunity of its specialist host locust

**DOI:** 10.1101/2020.06.13.149757

**Authors:** Zeyuan Jiang, Petros Ligoxygakis, Yuxian Xia

## Abstract

Conidial hydrophobins in fungal pathogens of plants^1,2^, insects^3,4^, and humans^5,6^ are required for fungal attachment and are associated with high virulence. They are believed to contribute to the pathogenesis of infection by preventing immune recognition^5,6^. Here, we refute this generalisation offering a more nuanced analysis. We show that MacHYD3, a hydrophobin located on the conidial surface of the specialist entomopathogenic fungus *Metarhizium acridum*, activates specifically the humoral and cellular immunity of its own host insect, *Locusta migratoria manilensis* (Meyen) but not that of other non-host insects. When topically applied to the cuticle, purified MacHYD3 improved the resistance of locusts to both specialist and generalist fungal pathogens but had no effect on the fungal resistance of other insects, including *Spodoptera frugiperda* and *Galleria mellonella*. Hydrophobins extracted from the generalist fungal pathogens *M. anisopliae* and *Beauveria bassiana* had no effect on the resistance of locusts to fungal infection. Thus, the host locust has evolved to recognize the conidial hydrophobin of its specialist fungal pathogen, whereas conidial hydrophobins from generalist fungi are able to evade recognition. Our results distinguish the immunogenic potential of conidial hydrophobins between specialist and generalist fungi.

## 1. Introduction

Fungal pathogens cause serious human, animal, and plant diseases and have numerous effects on human life^7^. Fungi are responsible for a wide array of superficial and disseminated (occasionally life-threatening) infections in humans, including *Candida albicans*, *Aspergillus fumigatus*, and *Cryptococcus neoformans*. Most human fungal pathogens are ubiquitous in the environment and humans are exposed to them by inhaling their spores^8^. But also in many others contexts fungi are the most ubiquitous pathogens. For example, they reduce populations of plants and insects, cause many of the most serious crop diseases^9^, and regulate insect populations in nature^10^. Therefore, fungi have great potential utility in controlling pest insects and weeds^11^.

To combat fungal diseases in agriculture and to develop fungal pesticides, extensive efforts have been made to clarify the molecular interactions between fungal pathogens and their hosts. Conidial attachment is the first crucial step in fungal infection and is modulated by hydrophobins^4,9^. Hydrophobins are small (molecular mass < 20 kDa), secreted hydrophobic proteins ubiquitously produced by filamentous fungi^4,12,13^. Although very diverse in their amino acid sequences, the hydrophobins constitute a closely related group of morphogenetic proteins^1^. Fungi can express two classes of hydrophobin, which play different roles in fungal growth, cell-surface properties, and development^4^. In the entomopathogenic fungus *Metarhizium brunneum*, *HYD1* and *HYD3* encode class I hydrophobins, and *HYD2* encodes a class II hydrophobin. Hydrophobins play roles in both fungal pathogenicity and infection-related development. The deletion of the three hydrophobin genes from the entomopathogenic fungi *M. brunneum* reduced its virulence^4^. In *Beauveria bassiana*, the inactivation of *HYD1* reduced spore hydrophobicity and fungal virulence, but *hyd2* mutants show reduced surface hydrophobicity with no effect on virulence^3^. In the rice blast fungus *Magnaporthe grisea*, the targeted replacement of *MPG1,* a gene encoding a fungal hydrophobin, produced mutants with reduced pathogenicity^14^. The hydrophobin MHP1, has essential roles in surface hydrophobicity and infection-related fungal development, and is required for the pathogenicity of *Magnaporthe grisea*^15^. Overall, the hydrophobins of fungal pathogens of plants and insects have key functions in pathogenicity, involving conidial attachment^3,14^, fungal germination^3^, the appressorium^4,14^, and host colonization^15^.

In addition to regulating infection-related development, hydrophobins can modulate the immune recognition and phagocytosis of human fungal pathogens. The conidial cell wall of *A. fumigatus* makes the airborne conidia resistant to the host’s immune response^6^. Mutants with impaired hydrophobins reportedly have a reduced hydrophobin layer^16^, and a lack of hydrophobins make conidia more susceptible to killing by alveolar macrophages^17^. However, hydrophobin itself cannot activate the host immune response^6^. Another study showed that the hydrophobins of spores mask the spores from the dectin-1- and dectin-2-dependent responses and enhances fungal survival^18^. These findings indicate that the hydrophobins of human pathogenic fungi prevent conidial recognition by the host immune system. In the interaction between the specialist pathogenic fungus *M. acridum* and its host insect, the humoral immunity of the locust responds quickly to the conidia on the fungal cuticle^19^, and the locust can detect the β-1,3-glucan of the fungal pathogen before its penetration, defending against infection via the Toll signalling pathway^20^. In addition to Toll, acute phase reactions in insects include the induction of phenoloxidase cascade, an insect-specific reaction akin to complement activation of cellular immunity with the rapid engagement of phagocytes to engulf the invading pathogen^21^. However, the effects of the hydrophobins of entomopathogenic fungi on the immune response of host insects have not been investigated. In this study, the effects of conidial hydrophobins of entomopathogenic fungi on the humoral and cellular immune responses of a host insect were evaluated, specifically the interaction between the conidial hydrophobin of *M. acridum* and the immune system of its specialist host *L. migratoria.* A hydrophobin of the specialist *M. acridum*, MacHYD3, activated specifically the humoral and cellular immunity of its own host insect, *L. migratoria* but not that of other insects, and improved the resistance of locusts to both specialist and generalist fungal pathogens but had no effect on the fungal resistance of other insects. However, hydrophobins extracted from the generalist fungal pathogens *M. anisopliae* and *Beauveria bassiana* had no effect on the resistance of locusts to fungal infection. The results showed that HYD3 of the the specialist *M. acridum* specifically induces locust immunity.

## 2. Materials and Methods

### 2.1 *Locusta migratoria manilensis* (Meyen) and fungal strains

*Locusta migratoria manilensis* (Meyen) is maintained in our laboratory under crowded conditions, as previously described^22^. Briefly, *L. migratoria* was maintained at 30 °C and 75% relative humidity with a 12:12 h light:dark photoperiod. The conidia of *M. anisopliae* var. *acridum* strain CQMa102 used in this study were provided by the Genetic Engineering Research Centre School of Life Science at Chongqing University, Chongqing, China, and were cultured as previously described^23^. Briefly, in a two-phase fermentation process, mycelia were first produced in a liquid fermentation reactor and then used to inoculate rice autoclaved with 40%-50% water in compound plastic bags (which permitted gas exchange but prevented microbial contamination). After 15 days, the conidia were harvested, dried, and used for hydrophobin extraction. For the bioassay, the conidia were cultured on 1/4 Sabouraud dextrose agar with 1% yeast extract (SDAY) for 2 weeks.

### 2.2 Surface hydrophobin extraction and purification

Hydrophobin was extracted from the spore surface as described previously^6^. Briefly, dry conidia were incubated with 48% hydrofluoric acid for 72 h at 4 °C. The samples were centrifuged (9,000 × g, 10 min) and the supernatant was dried under N_2_. The dried material was reconstituted in H_2_O. Hydrophobin was purified as previously described, with some modifications^24^. Briefly, the solution was applied to a column of highly substituted Phenyl Sepharose® 6 Fast Flow (Pharmacia Biotech, New York, USA) equilibrated with 100 mM Tris/HCl (pH 7.5) containing 2 M ammonium sulfate. Most of the hydrophobin was eluted with water after a linear gradient of the equilibrium buffer to 20 mM Tris/HCl (pH 7.5). The hydrophobin-containing fractions was further purified with anion exchange fast-performance liquid chromatography (Q-Sepharose; Pharmacia Biotech) to separate the different forms of hydrophobin. The proteins were eluted with a linear gradient of 0-0.5 M NaCl in 20 mM Tris/HCl (pH 9.0). The final hydrophobin preparation was concentrated by ultrafiltration (YM1 membrane; Amicon) and the solvent changed to water with gel filtration (Biogel P6-DG; Bio-Rad, USA). An aliquot was subjected to SDS-PAGE (15% gel), visualized with Coomassie Brilliant Blue staining with standard protocols, and confirmed with mass spectrometry (MS).

### 2.3 *Locusta migratoria* treated topically with hydrophobin, and hemolymph collection

Hydrophobin (20 μg) was topically applied to the locusts. The control groups were treated topically with 20 μg of bovine serum albumin (BSA). The locusts were then housed in groups of 10 individuals and fed maize leaves. Hemolymph was collected as described previously^25^. Briefly, Hemolymph was collected from the arthrodial membrane of the hind leg of each locust at 0.5, 1, 2, and 3 h after the topical application of hydrophobin or BSA. The arthrodial membrane of the hind leg of each locust was first swabbed with 70% ethanol, and then pierced with a sterile needle. After all the hemolymph was flushed from each locust, it was collected immediately and mixed with an equal volume of 0.5% sodium citrate to prevent coagulation.

### 2.4 Total hemocyte counts and phagocyte counts

Hemolymph samples (10 μL) were loaded onto a hemocytometer and the total numbers of cells and phagocytes were estimated under a compound microscope at 10× magnification. A Wright-Giemsa staining assay was used to count the phagocytes as described previously^24^. All experiments were repeated at least three times. The following equation was used to estimate the percentage of phagocytes:

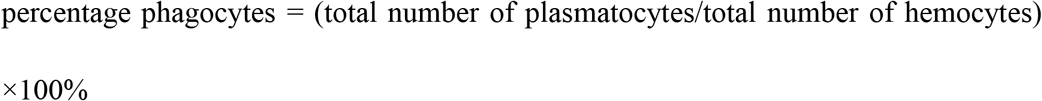

### 2.5 Extracellular phenoloxidase (PO) activity

Twenty fifth-instar nymphs of *L. migratoria* were topically inoculated with 5 μL of a paraffin oil suspension of *M. acridum* conidia (1 × 10^8^ conidia/mL) or 20 μg of hydrophobin from the spores of *M. acridum* and incubated for 1 h. The blood was then extracted from the locusts. Each blood sample was centrifuged at 30 × g for 10 min at 4 °C to remove the blood cells, and the PO activity was measured as described previously, with some modifications^26^. One unit of PO activity was defined as the change in absorbance at a wavelength of 490 nm (ΔA_490_) after 60 min = 0.001.

### 2.6 Immune-related gene transcription

Thirty fifth-instar of *L. migratoria* were topically treated with 20 μg of hydrophobin, 5 μL of *M. acridum* conidial suspension (1 × 10^8^ conidia/mL) or 20 μg of BSA. The fat bodies were collected after treatment with hydrophobin for 0.5, 1, 2, or 3 h. The total RNA was extracted with an Ultrapure RNA kit (CWbiotech) and the complementary DNA (cDNA) was synthesized. Quantitative real-time PCR (qPCR), and the data analysis were performed as previously described^27^. All the primers used for qPCR in this study are listed in Supplementary Table 1.

### 2.7 Subcellular localization of MacHYD3

The subcellular location of MacHYD3 in *M. acridum* was determined. Conidia of *M. acridum* were harvested from 1/4 SDAY plates after growth for 15 days at 28 °C. The conidia were incubated with a primary anti-MacHYD3 antibody overnight at 37 °C, washed five times with 1 × phosphate-buffered saline (PBS), incubated with a fluorescein isothiocyanate (FITC)-conjugated goat anti-rabbit IgG secondary antibody for 4 h at 37 °C, and washed five times with 1 × PBS. The subcellular location of MacHYD3 was determined as the distribution of green fluorescence, detected with fluorescence microscopy (Nikon Y-TV55, Tokyo, Japan). The predicted antigenic sequence at the N-terminus of MacHYD3 (amino acids 73-87: AIVPFGVKDGTGIRC) was synthesized commercially and used to raise antibodies in New England white rabbits entrust Cohesion Biosciences (UK). The control group was vaccinated with preimmune serum instead of the anti-MacHYD3.

### 2.8 Gene synthesis, subcloning, and expression

The complete gene sequence of MacHYD3 was determined, synthesized, and subcloned into the target vector pET-32a(+) for expression in *Escherichia coli*. The expected molecular weight of the expressed protein was ~25 kDa. cloning strategy: ATG-Trx-His tag-Kpnl-TEV protease site-protein-stop codon-*Hin*dIII. Competent *E. coli* BL21(DE3) cells stored at −80 °C were thawed on ice. The plasmid DNA (100 ng) was added to the *E. coli* BL21(DE3) cells and mixed gently. The tube was incubated on ice for 30 min and then heat shocked at 42 °C for ~90 s without shaking. Luria-Bertani (LB) medium (100 μL) at room temperature was added, and the tube was incubated with shaking at ~200 rpm for 60 min at 37 °C. The sample was spread on an LB agar plate containing 100 μg/mL ampicillin, and incubated upside down at 37 °C overnight. Two single well-isolated colonies were picked and used to inoculate 4 mL of LB broth containing 100 μg/mL ampicillin. The cells were incubated at 37 °C with shaking at 200 rpm. When A_600_ was 0.6~0.8, isopropyl β-D-1-thiogalactopyranoside (IPTG) was added to one tube at a final concentration of 0.5 mM IPTG to induce protein expression and the cells were incubated at 37 °C for 4 h. Another one without IPTG was used as the negative control. Protein expression and solubility were detected with SDS-PAGE.

### 2.9 Bioassays

Bioassays (topical inoculation and injection) were conducted with fifth-instar nymphs of *L. migratoria,* as described previously^28^. For the topical inoculation, the head-thorax junction of each locust was dipped into 5 μL of a paraffin oil suspension containing 1 × 10^7^ conidia/mL or the head-thorax junction of each locust was dipped into 40 μg of hydrophobin for 1 h and then into 5 μL of a paraffin oil suspension containing 1 × 10^7^ conidia/mL. The control locusts were treated with 5 μL of paraffin oil only. For the injections, 5 μL of an aqueous suspension containing 1 × 10^7^ conidia/mL or 5 μL of an aqueous suspension containing 1 × 10^7^ conidia/mL together with 20 μg of hydrophobin was injected into the hemocoel through the second or third abdominal segment of each locust. The control locusts were treated with 5 μL of sterile distilled water. Three replicates of each treatment were performed with 30 insects, and the experiment was repeated three times. Second-instar nymphs of *S. frugiperda* were dipped into 0.5 μL of a paraffin oil suspension containing 1 × 10^8^ conidia/mL on backside or each *S. frugiperda* was dipped into 5 μg of hydrophobin for 1 h and then into 5 μL of a paraffin oil suspension containing 1 × 10^8^ conidia/mL. The control were treated with 0.5 μL of paraffin oil only. Larvae of *G. mellonella* were inoculated by immersion in the spore suspension of *M. anisopliae* (1 × 10^8^ conidia/mL) for 20 s or dipped in 5 μg of MacHYD3 for 1 h and then inoculated by immersion in the spore suspension of *M. anisopliae* (1 × 10^8^ conidia/mL) for 20 s. Mortality was monitored at 12 h intervals.

### 2.10 Nodule counts

To detect the formation of nodules, 5 μL of an aqueous suspension containing 1 × 10^8^ conidia/mL or 5 μL of aqueous suspension containing 1 × 10^8^ conidia/mL together with 20 μg of MacHYD3 was injected into the hemocoel of 10 fifth-instar nymphs of *L. migratoria*. The number of nodules was calculated as previously described, with some modifications^29^. Briefly, the locusts were collected 12 h after injection. A mid-dorsal cut was made along the full length of the body. The gut and fat bodies were removed to expose the inner dorsal surface of the body wall. The nodules were counted routinely in all abdominal segments under a dissecting microscope. All experiments were repeated three times.

### 2.11 Statistical analysis

All values are expressed as means ± standard deviations. Statistical analyses were performed with the GraphPad Prism 7 software. The data from the survival experiments were analysed with a log-rank (Mantel-Cox) test. P values < 0.05 were considered statistically significant. An unpaired *t* test (two-tailed) and one-way analysis of variance (ANOVA) with a Tukey post hoc test for multiple comparisons were used to analyse the total hemocyte counts and the gene expression data. All the figures were generated with the same program.

## 3. Results

### 3.1 Conidia of *M. acridum* activate the cellular and humoral immune responses of locusts before germination

Spatial and temporal transcriptomic analyses have shown that the humoral immunity of the locust responds quickly to the conidia of the specialist pathogen *M. acridum* on its cuticle^19^. To confirm that *M. acridum* spores activate both the cellular and humoral immune responses of the locust before their germination, fifth-instar *L. migratoria* were topically treated with a suspension of *M. acridum* spores. The numbers of hemocytes and phagocytes, the expression of humoral-immunity-related genes of *L. migratoria* (*Lmspätzle*, *LmMyD88*, and *LmPPO11*), and phenoloxidase (PO) activity were measured. The total number of hemocytes was significantly reduced after treatment at 1-3 h post-inoculation (*P* = 0.022–0.0002; Fig. 1a, b), and the percentage of phagocytes was significantly higher after treatment than in the control groups at 1 h post-inoculation (*P* = 0.04; Fig. 1c). Transcription of two Toll pathway signalling components namely, the extracellular ligand *Lmspätzle* and intracellular receptor-adaptor complex mediator*LmMyD88*, as well as the pro-phenoloxidase gene *LmPPO11* was significantly increased at 1, 2, and 3 h post-inoculation (*P* = 0.0002, 0.0008, and 0.0007, respectively; Fig. 1d). Similarly, PO activity was significantly higher in the hemolymph of challenged *L. migratoria* than in that of the control group at 1 h post-inoculation (*P* = 0.0114; Fig 1e). These results demonstrate that the conidia of *M. acridum* activate the cellular and humoral immune responses of *L. migratoria* in the very early stage of infection, before germination.

**Figure 1.**
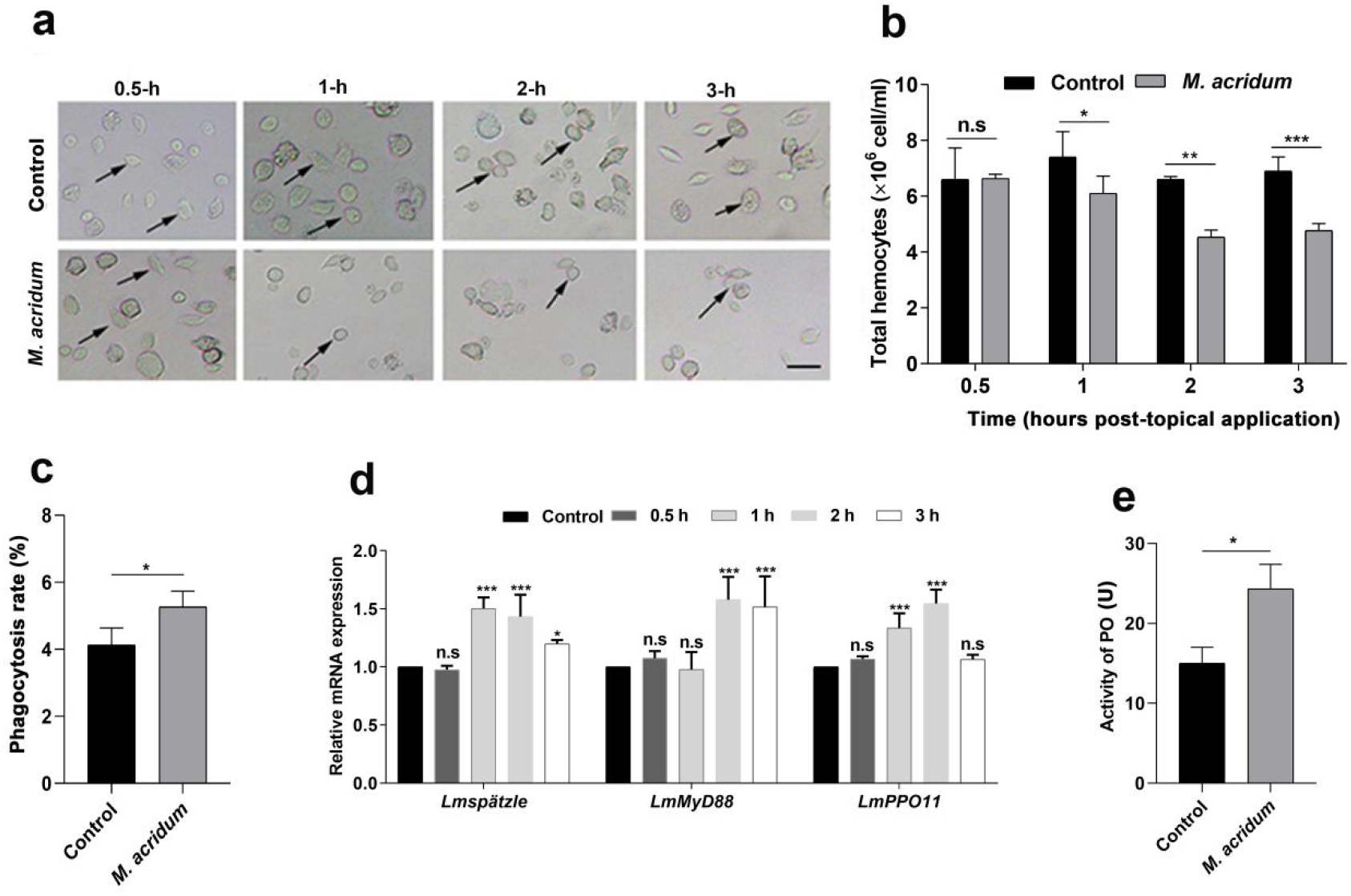
*Metarhizium acridum* activates the immune response of *Locusta migratoria*. **(a)** Representative images of *L. migratoria* hemocytes after topical application of 5 μl of spore suspension of *M. acridum* (1 × 10^8^ conidia/ml) relative to the control. Scale bar = 20 μm. Arrows indicate hemocytes. **(b)** Total hemocytes counts of *L. migratoria* 0.5, 1, 2, or 3 h after topical application of 5 μl of spore suspension of *M. acridum* (1 × 10^8^ conidia/ml). **(c)** Hemocytes engaged in phagocytosis 1 h after topical application of 5 μl of spore suspension of *M. acridum* (1 × 10^8^ conidia/ml. **(d)** Immunity-related gene expression in fat bodies of *L. migratoria* 0.5, 1, 2, or 3 h after topical application of 5 μl of spore suspension of *M. acridum* (1 × 10^8^ conidia/ml), n = 30 per group. **(e)** Phenoloxidase (PO) activity of *L. migratoria* 1 h after topical application of 5 μl of spore suspension of *M. acridum* (1 × 10^8^ conidia/ml). One unit of PO activity was defined as ΔA_490_ = 0.001 after 60 min, n=10 for each group except **(d)**. Control groups were treated with topical applications of 5 μl of paraffin oil **(a-e)**. n.s. *P* > 0.05, \**P* < 0.05, \*\**P* < 0.01, \*\*\**P* < 0.001. Error bars represent standard deviations of the means.

### 3.2 The hydrophobin of *M. acridum*, MacHYD3, is located on the conidial surface

Because the conidia of *M. acridum* activate the locust immune response before germination, and hydrophobin is located in the outermost layer of the conidial surface^30^, we speculated that the conidial hydrophobin of *M. acridum* plays a role in activating locust immunity. To test this hypothesis, the hydrophobin of *M. acridum* was extracted and purified to homogeneity with hydrophobic interaction chromatography, followed by anion exchange chromatography (Fig. 2a, b). Analysis of the amino acid sequence of the purified protein with mass spectrometry (MS) (Fig. 2c) showed that it was a hydrophobic protein (Fig. 2d) with eight conserved cysteines and a signal peptide (Fig. 2c,e). Alignment of the amino acid sequence of the purified protein with the genomic database of the *M. acridum* strain (CQMa102), other *Metarhizium* spp., and other ascomycetes showed that it was hydrophobin 3 (MacHYD3). The amino acid sequences of the hydrophobins were not conserved across different fungal species, or even within the same fungal strain (Fig. 2e). The phylogenetic relationships and classification showed that MacHYD3 is a class I hydrophobin (Fig. 2f), and a homologue of a previously reported *M. brunneum* hydrophobin^4^. To confirm the location of MacHYD3 on the conidia, an antibody directed against MacHYD3 was raised as described in the Materials and Methods. An immunohistochemical analysis showed that MacHYD3 was located on the surfaces of *M. acridum* conidia (Fig. 2g). Together, these results demonstrate that MacHYD3 is a class I hydrophobin located on the surface of the *M. acridum* conidium.

**Figure 2.**
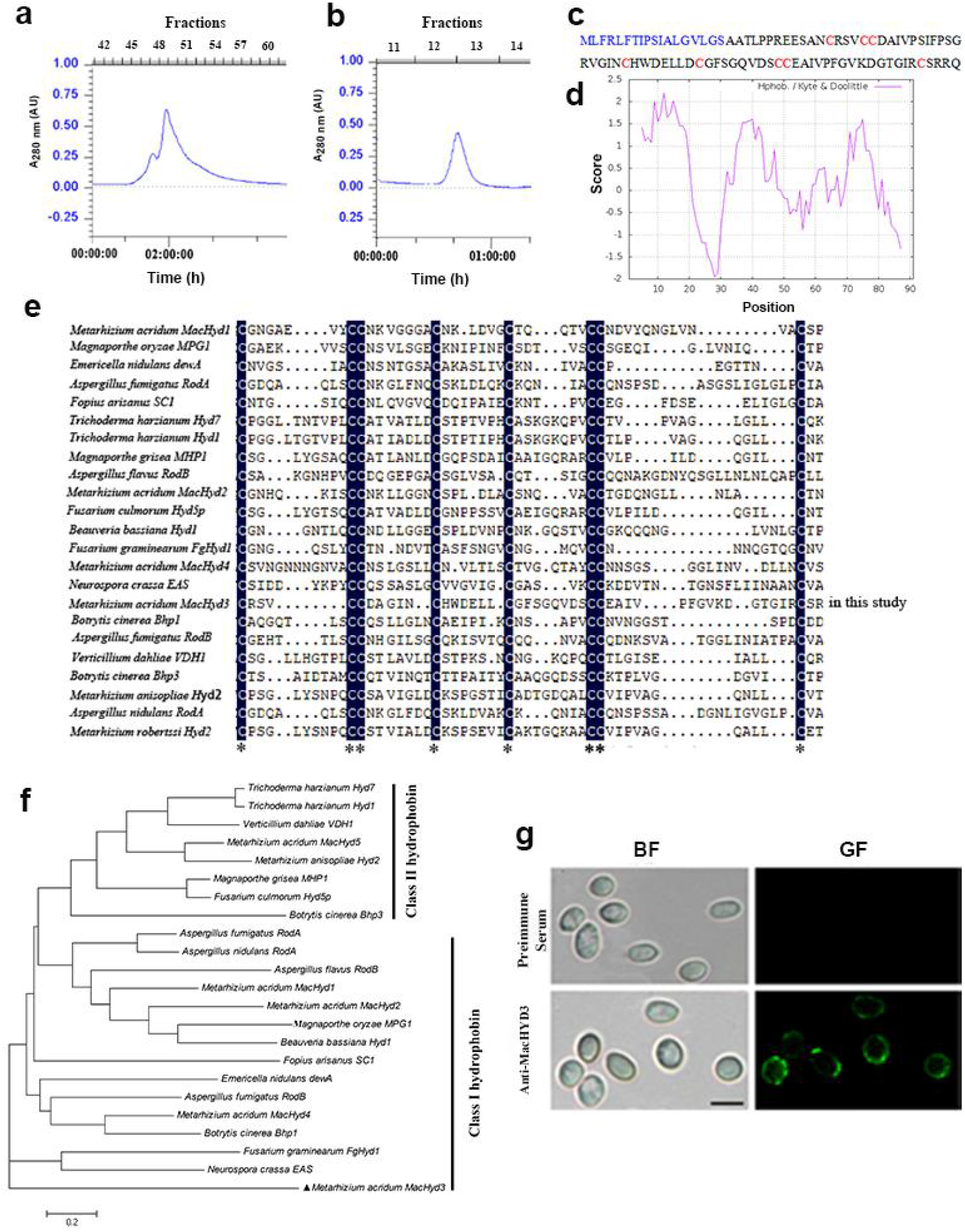
Analysis of hydrophobin MacDYD3, extracted from *Metarhizium acridum*. HPLC chromatograms of primary extracted hydrophobin. Hydrophobin after hydrophobic interaction chromatography **(a)** and anion exchange chromatography **(b)**. **(c)** Sequence of hydrophobin from *M. acridum*. (Signal peptide (in blue), eight conserved cysteines (in red). (Signal peptide prediction-http://www.cbs.dtu.dk/services/SignalP/). **(d)** ProtScale of hydrophobin (https://web.expasy.org/protscale/). **(e)** Alignment of the hydrophobin sequences from *L. acridum* genomes. Asterisk (*), denotes identity. **(f)** Phylogenetic relationships of MacHYD3 (black triangles) with hydrophobins from *M. acridum* strain CQMa102 and other *Metarhizium* spp. or other ascomycetes. The neighbor-joining model was constructed with the MEGA 5 software. The robustness of the generated tree was determined with 1,000 bootstrap replicates. The fungal proteins used were: *M. acridum* MacHYD1 (XP_007813670); *M. acridum* MacHYD2 (XP_007810716); *M. acridum* MacHYD3(XP_007809299); *M. acridum* MacHYD4 (XP_007811435); *M. acridum* MacHYD5 (XP_007815847); *M. anisopliae* Hyd2 (ADP37438); *Beauveria bassiana* Hyd1 (XP_008599918); *Botrytis cinerea* Bhp1 (BC1G_15273); *B. cinerea* Bhp3 (BC1G_01012); *Trichoderma harzianum* Hyd7 (AWT58102); *Trichoderma harzianum* Hyd1 (ANU06237); *B. bassiana* Hyd1 (XP_008599918); *Verticillium dahliae* VDH1 (AAY89101.1); *Magnaporthe grisea* MPG1 (P52751); *M. grisea* MHP1 (AAD18059); *Fusarium culmorum* FcHyd5p (ABE27986.1); *Neurospora crassa* EAS (EAA34064.1); *Claviceps purpurea* CPPH1(CAD10781.1); *Aspergillus nidulans* RodA (AAA33321.1); *A. flavus* RodB (XP_002375446); *A. nidulans* DewA (AAC13762.1)*; A. fumigatus* RodA (AAB60712.1), and *A. fumigatus* RodB (EAL91055.1). The corresponding accession numbers were obtained from the NCBI database (http://www.ncbi.nlm.nih.gov/). **(g)** Subcellular localization of MacHYD3. **Above**, Localization of MacHYD3 with pre-immune serum and a FITC-conjugated goat anti-rabbit IgG secondary antibody. **Below**, Localization of MacHYD3 with anti-MacHYD3 primary antibody and a FITC-conjugated goat anti-rabbit IgG secondary antibody. BF, bright field; GF, green fluorescence. Scale bar = 5 μm.

### 3.3 Conidial hydrophobin of *M. acridum*, MacHYD3, activates locust immunity but conidial hydrophobins from generalist species do not

To test whether MacHYD3 mediates the immunity-based interactions between *M. acridum* and *L. migratoria*, we measured the effects of MacHYD3 purified from conidial preparations as well as recombinant MacHYD3 (rMacHYD3) on the immune response of *L. migratoria*. MacHYD3 or rMacHYD3 were topically applied to the head-thorax junction of *L. migratoria*. To monitor host defence, we measured the total number of hemocytes, the proportion of phagocytes among the hemocytes, the PO activity in the hemolymph, and the expression of immune-related genes (*Lmspätzle*, *LmMyD88*, and *LmPPO11*) in the fat body at 0.5, 1, 2, and 3 h after application (described in Materials and Methods). Compared with the control group, the total number of *L. migratoria* hemocytes was significantly reduced at 1 h (*P* = 0.0009; Fig. 3a, b), whereas the proportion of phagocytes (*P* = 0.0144; Fig. 3c) and PO activity (*P* = 0.0018; Fig. 3d) were significantly elevated at 1 h after the topical application of MacHYD3. The expression of *Lmspätzle*, *LmMyD88*, and *LmPPO11* was significantly upregulated at 1 h after the topical application of MacHYD3 (*P* = 0.0151, 0.0013, and 0.0021, respectively; Fig. 3e). Similarly, rMacHYD3 reduced the total number of hemocytes at 1 h (*P* = 0.0007; Fig. 3b) and increased PO activity in the hemolymph (*P* = 0.0049; Fig. 3d). These data suggest that conidial MacHYD3 activates both the cellular and humoral innate immune responses of *L. migratoria,* as do *M. acridum* conidia in the early stage of infection after conidial attachment.

**Figure 3.**
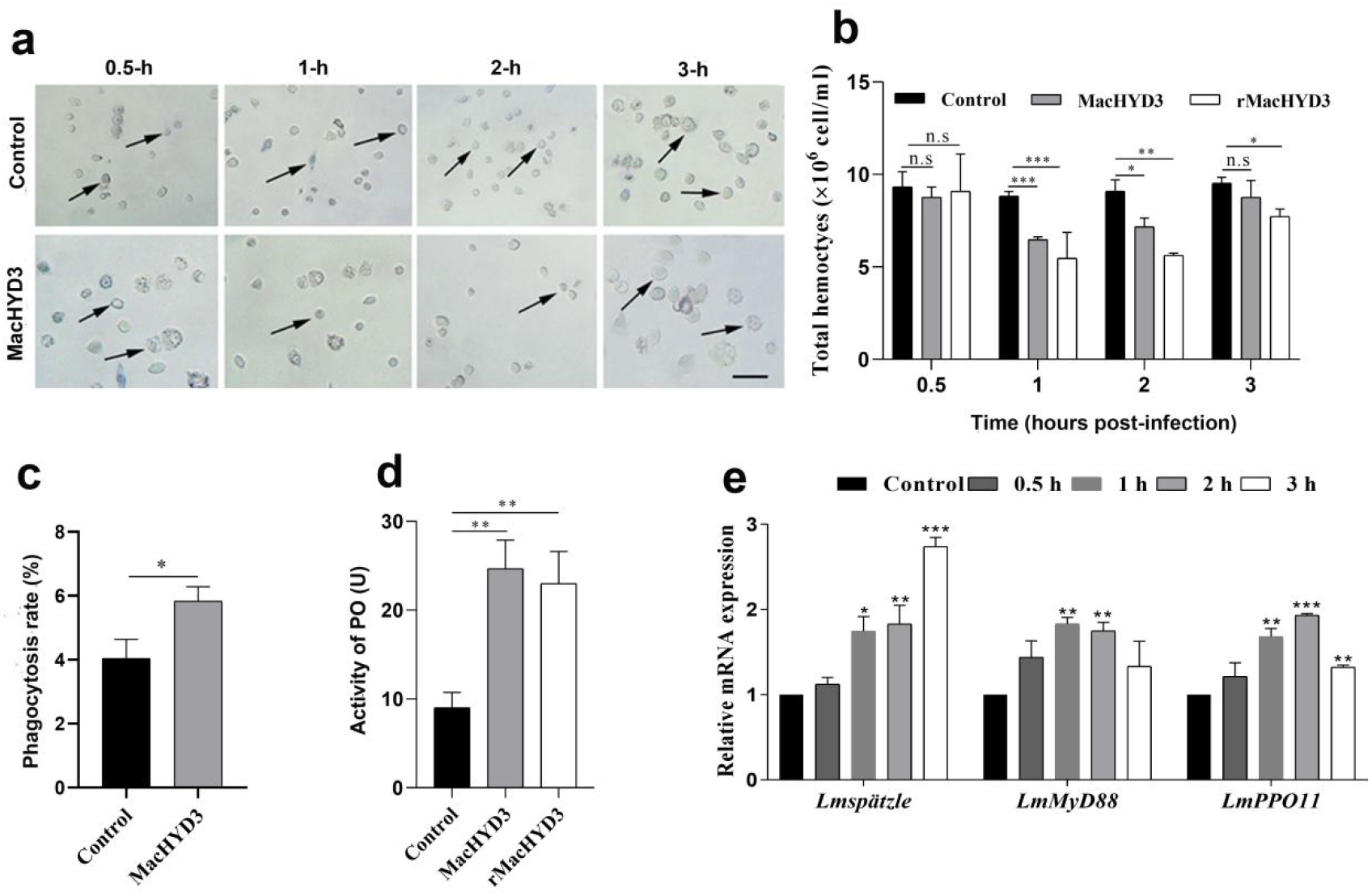
Hydrophobin of *Metarhizium acridum* activates the innate immune response of *Locusta migratoria*. **(a)** Total hemocyte preparation of *L. migratoria* after topical application of 20 μg of hydrophobin of *M. acridum* (MacHYD3). Scale bar = 20 μm. Arrows indicate hemocytes. **(b)** Total hemocyte counts of *L. migratoria* 0.5, 1, 2, or 3 h after topical application of 20 μg of hydrophobin of *M. acridum* (MacHYD3) of rMacHYD3 to the head-thorax junction. **(c)** Plasmatocytes of *L. migratoria* 1 h after topical application of 20 μg of MacHYD3 to the head-thorax junction. **(d)** Phenoloxidase (PO) activity of *L. migratoria* 1 h after topical application of 20 μg of MacHYD3 or rMacHYD3 to the head-thorax junction. One unit of PO activity was defined as Δ_A490_ = 0.001 after 60 min. **(e)** Immunity-related gene expression in fat body of *L. migratoria* 0.5, 1, 2, or 3 h after topical application of 20 μg of MacHYD3 to the head-thorax junction, n = 30 per group. Control groups were treated with topical application of 20 μg of bovine serum albumin (BSA), n = 10 per group, except **(e)**. n.s. *P* > 0.05, \**P* < 0.05, \*\**P* < 0.01, \*\*\**P* < 0.001. Error bars represent standard deviations of the means.

To test whether conidial hydrophobins from generalist fungi activate *L. migratoria* immunity, the hydrophobins of *M. anisopliae* (MaaHYD) and *B. bassiana* (BbHYD) were extracted. Neither MaaHYD nor BbHYD had a significant effect on the total number of hemocytes at 0.5, 1, 2 or 3 h after topical application (Fig. S1a) or PO activity 1 h after topical application (Fig. S1b). These results show that both these hydrophobins failed to elicit an innate immune response in *L. migratoria*.

Together, these data demonstrate that the conidial hydrophobin of the specialist fungal pathogen *M. acridum*, MacHYD3, activates the cellular and humoral immune responses of its host locust.

### 3.4 Conidial hydrophobin of *M. acridum*, MacHYD3, improves the resistance of *L. migratori*a but not of other insects, to both specialist and generalist fungi

Since MacHYD3 induces locust immune responses, we tested if it could prime the insect’s immune system and thus improve its resistance to infection by both specialist and generalist fungi. To estimate the effects of MacHYD3 on the locust’s resistance to pathogenic fungi, bioassays were conducted by injecting MacHYD3 into locusts or topically loading MacHYD3 onto the head–thorax junction of locusts, and then inoculating them with 5 μL of conidial suspension (1 × 10^7^ conidia/mL) from the specialist strain *M. acridum* CQMa102. The locusts treated with MacHYD3 died much more slowly, with a significantly higher median time-to-lethality (LT_50_), compared to the non-treated controls (*P* = 0.0162; Fig. 4a, 4b). Locusts injected with MacHYD3 lived as long after inoculation with *M. acridum* CQMa102 conidia as the uninfected controls (Fig. 4c). Significantly more nodules were observed in the inner dorsal body walls of the MacHYD3-injected locusts at 12 h after inoculation with *M. acridum* CQMa102 conidia (*P* = 0.0344; Fig. 4d, 4e). To test the effects of MacHYD3 on the locust’s resistance to a generalist fungus, locusts were loaded with MacHYD3 at the head–thorax junction and then topically inoculated with 5 μL of a conidial suspension (1 × 10^7^ conidia/mL) of *M. anisopliae* CQMa421. The locusts treated with MacHYD3 died more slowly, with a LT_50_ statistically higher, than that of the infected but not MacHYD3-treated locusts (*P* = 0.0002; Fig. 4f, 4g).

**Figure 4.**
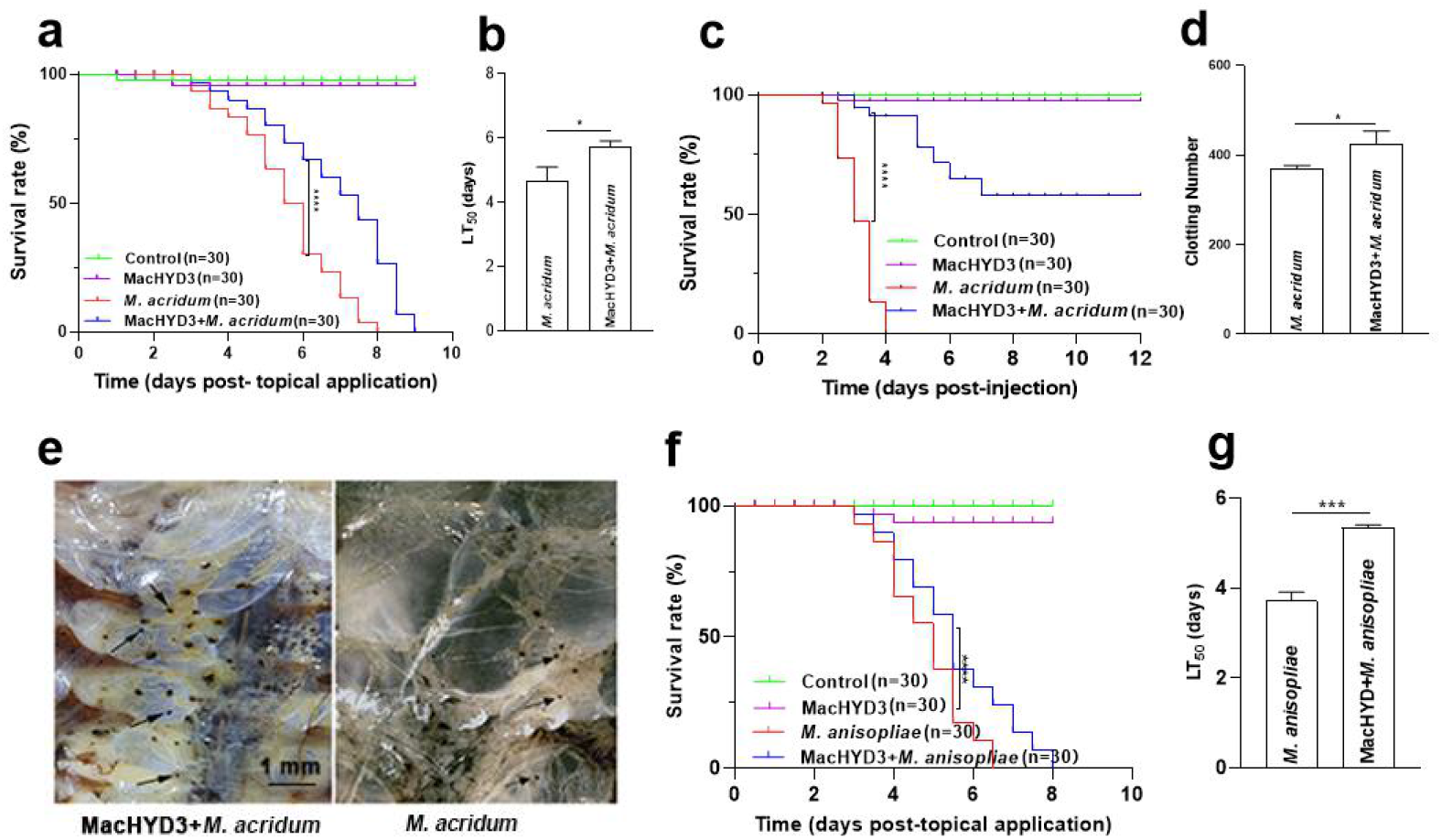
MacHYD3 primes the locust’s immune system for better survival after infection. **(a)** *Locusta migratoria* after topical application of 5 μl of conidial suspension of *Metarhizium acridum* (1 × 10^7^ conidia/ml) or/and 1 h after 40 μg of MacHYD3 was topical application. Control groups were treated with topical application of 5 μl of paraffin oil. **(b)** The median 50% lethality time (LT_50_) of *L. migratoria* after topical application of *M. acridum* or */*and MacHYD3. **(c)** *Locusta migratoria* injected with 5 μl of conidial suspension of *M. acridum* (1 × 10^7^ conidia/ml) or/and 20 μg of MacHYD3. Control groups were injected with 5 μl of sterile distilled water. Experiments were performed in triplicate. **(d)** Number of nodules on insect dorsal inner body walls 12 h after injection, \*\**P* < 0.01, n = 10. **(e)** Nodule formation in insect dorsal inner body walls at 12 h after injection. Arrows indicate typical nodules. Scale bar = 1 mm. **(f)** *Locusta migratoria* after topical application of 5 μl of conidial suspension of *M. anisopliae* (1 × 10^7^ conidia/ml), 40 μg MacHYD3 (MacHYD3), 5 μl 1×10^7^ conidia/mL condial suspension of *M. anisopliae* was topical application 1 h after 40 μg MacHYD3 was topical application or 5 μL of paraffin oil (Control). **(g)** The median 50% lethality time (LT_50_) of *L. migratoria* after topical application of *M. anisopliae* or *M. anisopliae* + MacHYD3. Data were analyzed with a log-rank test **(a,c,f)**. \*\*\*\**P* < 0.0001. Error bars represent standard deviations, \**P* < 0.05. \*\*\**P* < 0.001 **(b,d,g)**.

These results demonstrate that the hydrophobin MacHYD3 from a specialist fungal pathogen improved the resistance of its host insect, *L. migratoria*, to both specialist and generalist fungal pathogens. To test the efficacy of MacHYD3 in improving the resistance of non-host insects, MacHYD3 were loaded onto *S. frugiperda* and *G. mellonella* before they were inoculated with 5 μL of conidial suspension (1 × 10^7^ conidia/mL) of the generalist *M. anisopliae* strain CQMa421. There was no significant difference in locust survival after treatment with or without MacHYD3 (Fig. S2a, S2b, S2c and S2d). These data demonstrate that the conidial hydrophobin of *M. acridum*, MacHYD3 only improves the resistance of its host insect *migratoria* to both specialist and generalist fungi.

## 4. Discussion

As all external physical barriers, the insect cuticle is the first line of defence against fungal pathogens. In this study, our data demonstrate that the conidia of *M. acridum* activate both the cellular and humoral immune response of the locust before their germination. This is consistent with recent findings that the humoral immunity of the locust is activated soon after conidial attachment to the cuticle and during the germination of the specific pathogen *M. acridum*^19^. Therefore, the host immune response is activated much earlier during the invasion of the insect epidermis by the fungus than previously recognized. It had been thought that the lesions in the insect exoskeleton caused by the invading fungal cells were unlikely to elicit a detectable humoral immune response before the first layer of live cells in the epidermis had been breached by the fungus^31,32,33,34^. However, recent findings suggest that components of the fungal conidial surface elicit cellular and humoral immune responses in the host insect. β-1,3-Glucan is a component of the conidial surface and a pathogen-associated molecular pattern (PAMP), It localizes around the germinated conidia and at the germ-tube apex and infiltrates the hemocoel of the locust during the fungal germination stage, thus activating the Toll signalling pathway of the locust to defend it against fungal infection^20^. Our results here showed that the total number of hemocytes was significantly reduced after treatment at 1–3 h post-inoculation. As seen elsewhere this wasdue to the fact that hemocytes move and attach to the basement membranes of the epidermis beneath the host cuticle^35^. Moreover, the percentage of phagocytes and PO activity were significantly higher after treatment than in the control groups at 1 h post-inoculation. Taken together, these results indicate that the humoral and cellular immunity of the host insect responds very quickly to challenge by the fungal pathogen.

Along with β-1,3-Glucan, hydrophobins are also components of the conidial surface, located in the outermost layer^30^. Conidial hydrophobins play a crucial role in the attachment of fungal spores to the surface of the host insect^4,16^ and in infection-related development of fungi^3,4,14,15^. Nevertheless, they are considered “stealth” molecules that prevent induction of immune activity^6^. For example, *Aspergillus fumigatus* lacking the hydrophobin RodA is more susceptible to killing by alveolar macrophages^17^, but RodA itself does not activate the host immune system^6^, a result confirmed in a mouse infection model^36^. Our study confirms that the *M. acridum* hydrophobin MacHYD3 is located on the surface of the conidium. Unexpectedly however, we found that the conidial hydrophobin of *M. acridum*, MacHYD3, activates the cellular and humoral immune responses of its specialist host, the locust *L. migratoria*. In contrast, the conidial hydrophobins of generalist fungi failed to activate the innate immune response of *L. migratoria*. This result suggests that conidial hydrophobins from generalist fungi *M. anisopliae*, and *B. bassiana*, play similar roles in conidial masking and fungal immune evasion as the hydrophobin of *A. fumigatus*, RodA. Contradicting the idea that fungal hydrophobins act as a sheath to prevent the immune recognition of fungal conidia as in the generalist fungal species, the hydrophobin MacHYD3 from the fungus *M. acridum* activated both the cellular and humoral immunity of its specialist host. MacHYD3 also significantly improved the resistance of the host locust to both specialist and generalist fungi by priming its immune system. However, MacHYD3 failed to improve the resistance of non-host insects, *S. frugiperda* and *G. mellonella.* These data indicate that a conidial hydrophobin of the specialist fungal entomopathogen *M. acridum*, HYD3, specifically activates the cellular and humoral immune responses of its own host, *L. migratoria*. This specificity may be attributable to differences in the coevolution between generalist and specialist fungi with their hosts. Generalist fungi are often opportunistic pathogens, and their hosts may not have had sufficient opportunity to evolve mechanisms to recognize their conidial hydrophobins. However, during their long history of coevolution, the host of a specialist fungal species would have evolved an effective mechanism to recognize the conidial hydrophobins of that species. We hypothesise that such early detection of pathogens like *M. acridum*, is a prerequisite for an effective and successful defence as this is a time-sensitive response.

The two components of the cell wall of the *M. acridum* spore, hydrophobin (this study) and β-1,3-glucan (laminarin)^20^, activate the Toll signalling pathway when applied to the cuticle of *L. migratoria*. Because hydrophobin is located in the outermost layer of the conidium, whereas β-1,3-glucans are exposed when the conidium is germinating^6^, HYD3 of *M. acridum* should be the earliest immune inducer, whereas β-1,3-glucan may intensify this response during the germination stage. In generalist species, hydrophobin may act as a sheath, masking the PAMPs including β-1,3-glucan on the cell wall of the fungal conidium, to prevent immune recognition, so the host does not detect the generalist fungal conidium until germination. Therefore, the evidence presented in this study suggests that MacHYD3 of the specialist fungus *M. acridum* acts as the earliest PAMP for the immune system of its host *L. migratoria.*

An outstanding question concerns the molecular details of MacHYD3 recognition by the *L. migratoria* immune system as well as the mechanisms by which the recognition signal is transduced to Toll. Further studies are required to fully clarify these issues. Studies in *Drosophila* have shown that fungal proteases can act as “danger signals” to directly trigger the host proteolytic cascade leading to Spaetzle activation and Toll induction^37^. Whether hydrophobins can do the same or whether there is a specific *L. migratoria* recognition receptor remains to be identified. Uncovering the mechanism by which conidial hydrophobins are recognised could provide new strategies for the development of drugs to counter specialist fungi as well as the design of more effective fungal-based pesticides.

## Author Contributions

Design of the experiments, Y.X. and Z.J.; performance of the experiments, Z.J.; writing the manuscript, Y.X., P.L. and Z.J. All authors have read the final manuscript.

## Funding

This research was funded by grants from the National Key R & D Program of China (Project No. 2017YFD0201208) and the Natural Science Foundation of China (No. 31540089).

## Conflicts of Interest

The authors declare that they have no conflicts of interest.

---

**Supplementary Figure 1.**
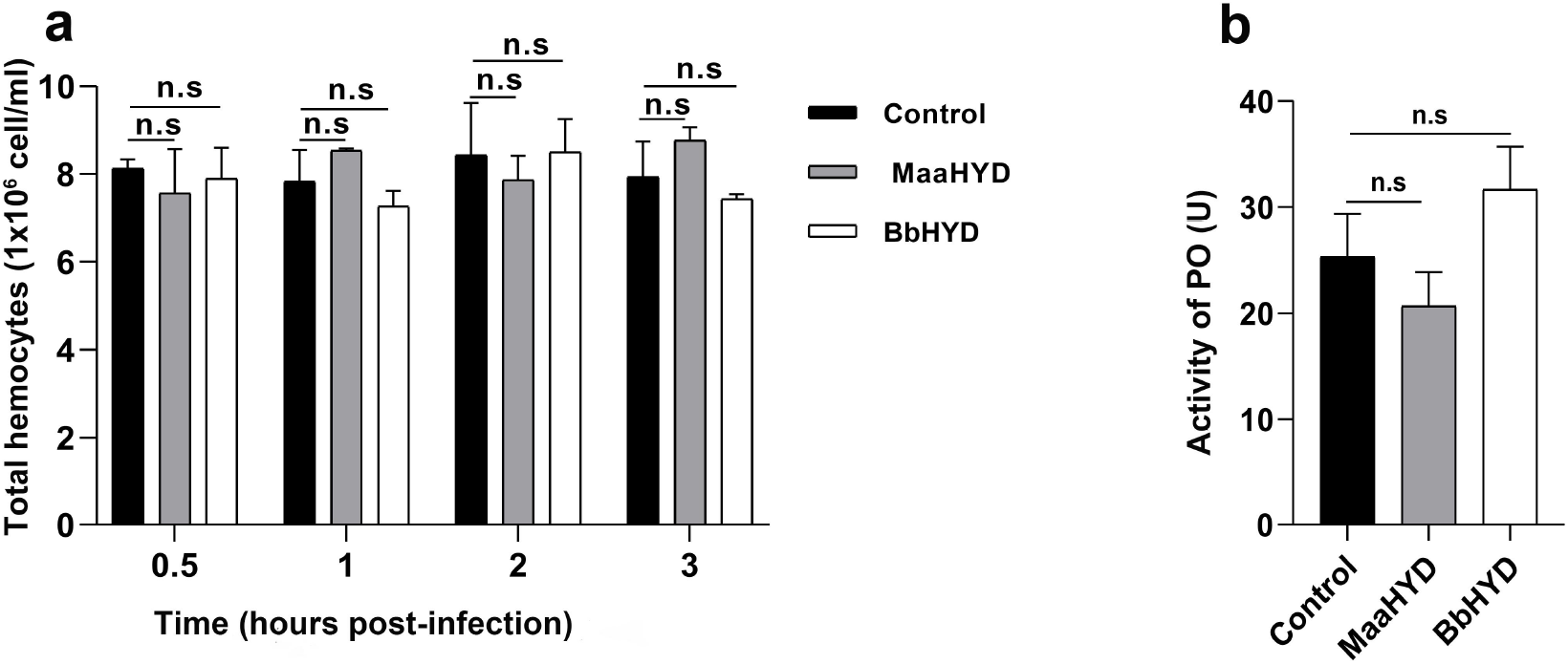
Hydrophobin from conidia of *Metarhizium anisopliae* (MaaHYD) and *Beauveria bassiana* (BbHYD) does not activate the innate immunity of *Locusta migratoria*. **(a)** Total number of hemocytes of *L. migratoria* 0.5, 1, 2, or 3 h after topical application of 20 μg of MaaHYD or BbHYD. **(b)** Phenoloxidase (PO) activity of *L. migratoria* 1 h after topical application of 20 μg of MaaHYD or BbHYD. One unit of PO activity was defined as ΔA_490_ = 0.001 after 60 min. Data are representative of at least two independent experiments, each with similar results. n.s., *P* > 0.05. Error bars represent standard deviations of the means.

**Supplementary Figure 2.**
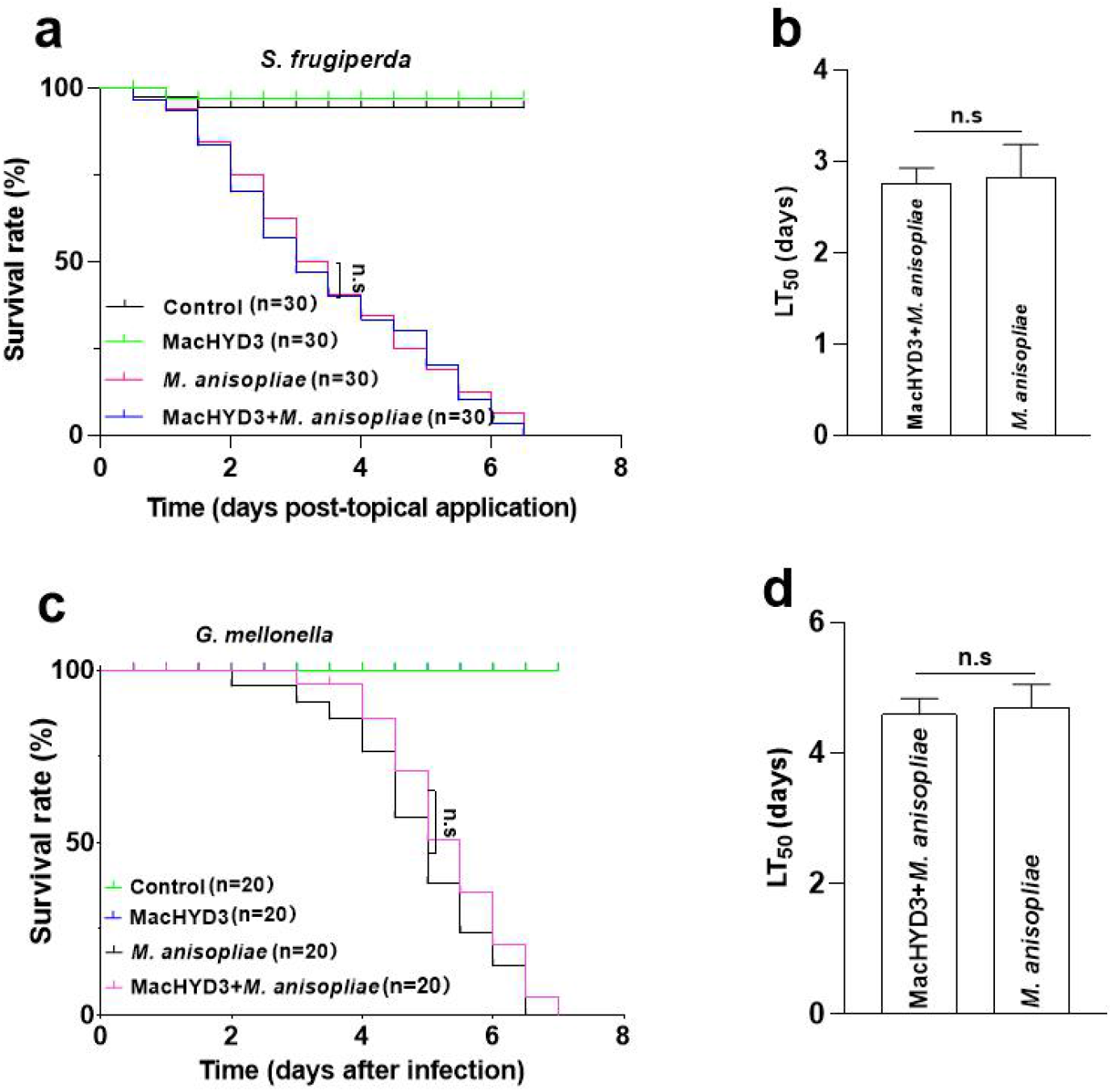
Survival after infection of non-host insects. **(a)** *Spodoptera frugiperda* after topical application of 0.5 μl of conidial suspension of *M. anisopliae* (1 × 10^8^ conidia/ml), 5 μg MacHYD3 (MacHYD3), 0.5 μl spore/mL condial suspension of *M. anisopliae* (1 × 10^8^ conidia/ml) was topical application 1 h after 5 μg MacHYD3 was topical application (MacHYD3+*M. anisopliae*), or 0.5 μl of paraffin oil (Control). **(b)** LT_50_ for *S. frugiperda* after topical application assay. **(c)** *Galleria mellonella* larvae were inoculated by immersion in the spore suspension for 20 s (1 × 10^7^ conidia/ml, *M. anisopliae*), larvae were immersed in spore suspension for 20 s (1 × 10^7^ conidia/ml, *M. anisopliae*) 1 h after 5 μg MacHYD3 was topical application (MacHYD3+*M. anisopliae*); larvae were immersed in sterile water (Control); or 5 μg MacHYD3 was topical application on backs of larvae (MacHYD3). **(d)** LT_50_ for *G. mellonella* larvae inoculated in the immersion assay. Data were analyzed with a log-rank test **(a,c)**, n.s., *P* > 0.05. Error bars represent standard deviations **(b,d)**, n.s., *P* > 0.05.

## Notes

### Competing Interest Statement

The authors have declared no competing interest.

