## Supplementary information for "HYD3, a conidial hydrophobin of the fungal entomopathogen *Metarhizium acridum* induces the immunity of its specialist host locust"

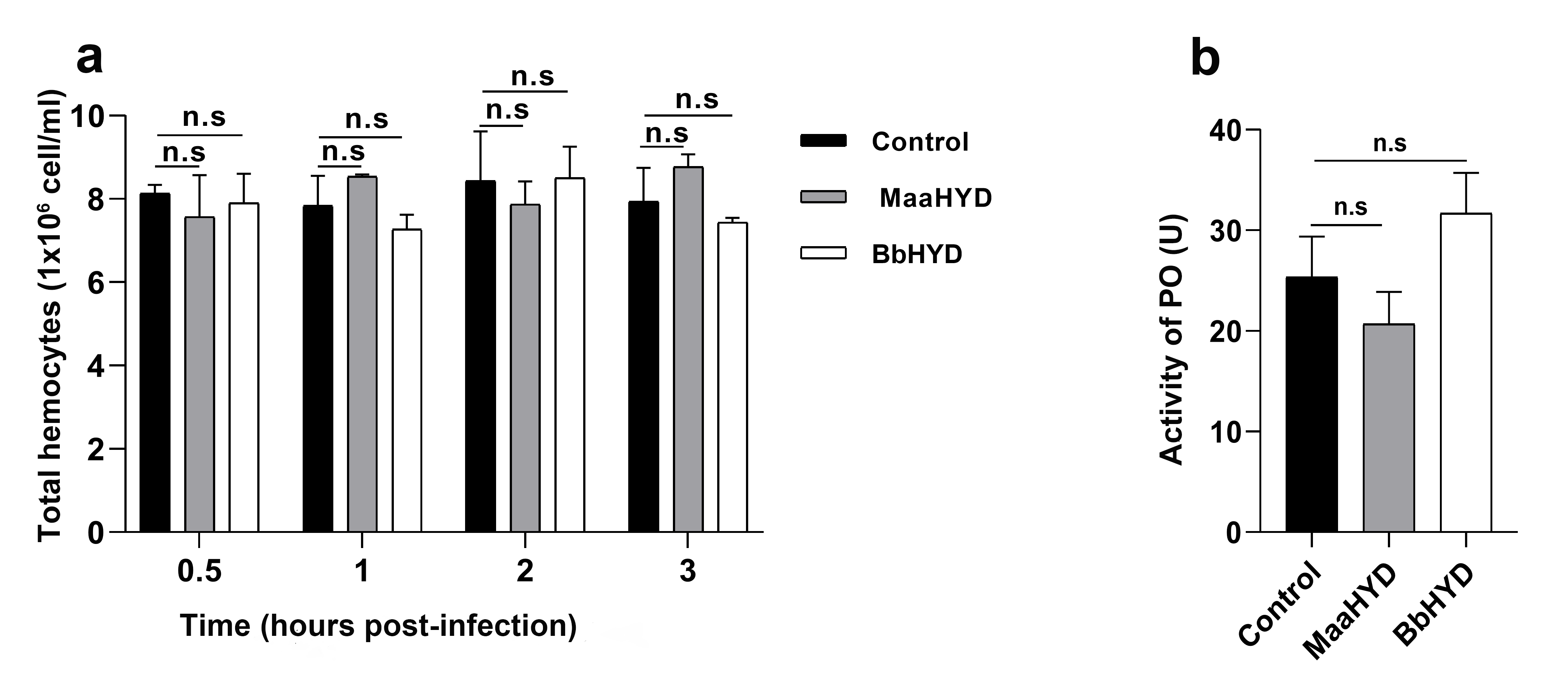
**Supplementary Fig. 1 Hydrophobin from conidia of *Metarhizium anisopliae* (MaaHYD) and *Beauveria bassiana* (BbHYD) does not activate the innate immunity of *Locusta [migratoria](C:/Users/%E8%92%8B%E6%B3%BD%E5%85%83/AppData/Local/youdao/DictBeta/Application/7.3.0.0817/resultui/dict/javascript:;)*.** **(a)** Total number of hemocytes of *L. [migratoria](C:/Users/%E8%92%8B%E6%B3%BD%E5%85%83/AppData/Local/youdao/DictBeta/Application/7.3.0.0817/resultui/dict/javascript:;)* 0.5, 1, 2, or 3 h after topical application of 20 μg of MaaHYD or BbHYD. **(b)** Phenoloxidase (PO) activity of *L. [migratoria](C:/Users/%E8%92%8B%E6%B3%BD%E5%85%83/AppData/Local/youdao/DictBeta/Application/7.3.0.0817/resultui/dict/javascript:;)* 1 h after topical application of 20 μg of MaaHYD or BbHYD. One unit of PO activity was defined as ΔA_490_ = 0.001 after 60 min. Data are representative of at least two independent experiments, each with similar results. n.s., *P* > 0.05. Error bars represent standard deviations of the means.

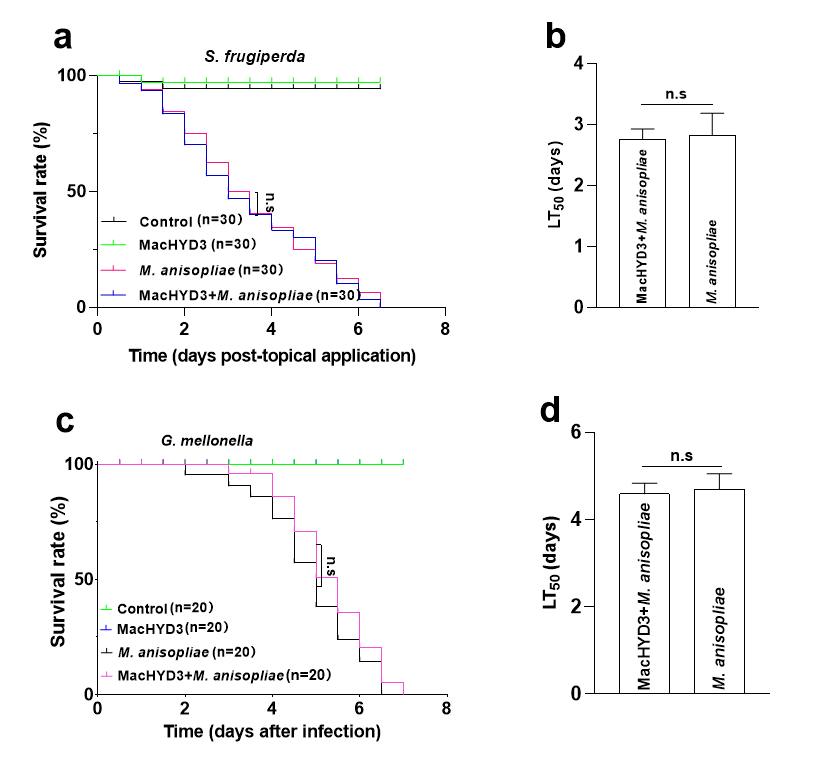

**Supplementary Fig. 2 Survival after infection of non-host insects**. **(a)** *Spodoptera frugiperda* after [topical](C:Usersèæ³½åAppDataLocalyoudaoDictBetaApplication7.3.0.0817resultuidict?keyword=topical) application of 0.5 µl of conidial suspension of *M. anisopliae* (1 × 10^8^ conidia/ml), 5 µg MacHYD3 (MacHYD3), 0.5 µl spore/mL condial suspension of *M. anisopliae* (1 × 10^8^ conidia/ml) was topical application 1 h after 5 µg MacHYD3 was topical application (MacHYD3+*M. anisopliae*), or 0.5 µl of paraffin oil (Control). **(b)** LT_50_ for *S. frugiperda* after [topical](C:Usersèæ³½åAppDataLocalyoudaoDictBetaApplication7.3.0.0817resultuidict?keyword=topical) application assay. **(c)** *Galleria mellonella* larvae were inoculated by immersion in the spore suspension for 20 s (1 × 10^7^ conidia/ml, *M. anisopliae*), larvae were immersed in spore suspension for 20 s (1 × 10^7^ conidia/ml, *M. anisopliae*) 1 h after 5 µg MacHYD3 was topical application (MacHYD3+*M. anisopliae*); larvae were immersed in sterile water (Control); or 5 µg MacHYD3 was topical application on backs of larvae (MacHYD3). **(d)** LT_50_ for *G. mellonella* larvae inoculated in the immersion assay. Data were analyzed with a log-rank test **(a,c)**, n.s., *P* > 0.05. Error bars represent standard deviations **(b,d)**, n.s., *P* > 0.05.

Supplementary table 1 primer sequences for qPCR

| gene | Forward Primer (5’>3’) | Reverse Primer (5’>3’) |
| --- | --- | --- |
| *OPP11* | GGAGAGAAATTCTCTTTACGG | CAGATTTAGCAGATAGATGGG |
| *spätzle* | AGCTTGTGGGTACGGAGAC | GGGCGATGAATAGATGAAAC |
| *MyD88* | CCTTTATGAGCGAGACTT | TCTTTGAGAGGACGACTA |
